# Putative signatures of genomic adaptation to Cuckoo brood parasitism in Reed Warbler hosts across Europe

**DOI:** 10.1101/2025.01.06.631542

**Authors:** William J. Smith, Katja Rönkä, Staffan Bensch, Nora M. Bergman, Rachael Y. Dudaniec, Fabrice Eroukhmanoff, Frode Fossøy, Bengt Hansson, Michał T. Jezierski, Edward Kluen, Petr Procházka, Camilla Lo Cascio Sætre, Bård G. Stokke, Rose Thorogood

## Abstract

Behaviour is a fundamental defence mechanism for animals facing enemies, yet its plasticity makes finding associated genomic evidence for coevolution challenging. As a solution, we apply landscape genomics methods more typically deployed with abiotic variables to a coevolutionary model system: Reed Warblers (*Acrocephalus scirpaceus*) and their Cuckoo (*Cuculus canorus*) brood parasites. Using continental sampling of genome-wide data from hosts breeding under varying brood parasitism rates, we reveal that gene flow varies among populations, and, controlling for variation due to environmental differences, identify genomic regions and putative genes that differ with exposure to brood parasitism. Therefore, our results suggest that variable host responses to selection are possible, even by highly mobile and connected populations, supporting long-held assumptions that behavioural defences may reflect genomic adaptation despite their plasticity. Further studies, including those linking genetic variation to individual-level behaviours involved in parental responses to brood parasitism, are required to confirm a genetic basis to specific coevolutionary adaptations, and to test for other biotic or abiotic variables potentially confounding with local parasitism rate.

## Introduction

Interactions between parasites and their hosts are often assumed to result in antagonistic coevolution, where counter-adaptation leads to reciprocal changes. In theory, one party will eventually drive the other to diversify or go extinct (1) but ecological variation across interacting species’ distributions is thought to maintain coevolution over the long-term (i.e. the Geographic Mosaic Theory of Coevolution, or GMTC (2)). Here, differential gene flow among populations of hosts and parasites is predicted to generate mosaics of varying selection (and responses) at the landscape level and reduce opportunities for either party to gain the upper hand. Testing this theory across a broad range of systems in the wild could lead to substantial advances in understanding links between evolutionary processes and macroecological patterns in a changing world (3). However, the key assumptions of the GMTC remain challenging to test without genomic evidence or common garden experiments (4). This is especially problematic when adaptations are behavioural and/or highly plastic (5) because the complex genetic basis of behavioural traits is usually only assumed (i.e. the “phenotypic gambit” (6)) and methods to scan genomes for such traits are still largely out of reach, particularly for mobile species where common garden approaches are unfeasible (7) and the level of experimental control possible is lower than for plant or invertebrate systems (e.g. 8). Consequently, geographic mosaics of coevolution among animal hosts and parasites in the wild remain largely descriptive with little evidence of underlying processes, even for measures of strength of selection or levels of gene flow (4). Behavioural resistance to parasitism is, however, a common strategy in nature (9) so it is necessary to find a method with which to detect genomic signals of selection and distinguish between behavioural plasticity and evolutionary change. Here we propose that genotype-environment analyses can provide a valuable first step in characterising the genetic basis of geographic mosaics of behavioural coevolution.

Avian brood parasitism is a textbook example of antagonistic coevolution. Cuckoos (*Cuculus canorus*) are highly virulent, laying host-specific eggs which hatch into chicks that evict the host’s offspring. In response, parent Reed Warblers (*Acrocephalus scirpaceus*) can protect their offspring by mobbing adult Cuckoos and/or ejecting their eggs. However, Cuckoos’ use of egg- and hawk-mimicry as a counter-adaptation ensures that there are potential costs to the Reed Warbler if a defence is launched inappropriately (10). Reed Warblers have therefore also evolved behavioural plasticity and rely on cues of local Cuckoo activity to fine-tune expression of defences according to local variation of parasitism risk (11). However, at sites where the Reed Warbler specific lineage (or ‘gens’) of Cuckoo does not occur (i.e. where selection is relaxed), the expression of defences varies from absent to persistent (11), suggesting that a geographic mosaic of variable strengths of selection (i.e. ‘hotspots’ and ‘coldspots’ (2)) is possible. Cuckoos and their hosts were given as a putative example of the GMTC in seminal publications (12), and the counter-adaptations of Cuckoos to defeat host defences can generate both genetic and phenotypic divergence (13, 14). Nevertheless, fundamental components of the GMTC for hosts are yet to be tested, and we can therefore still only take a phenotypic gambit in assuming that behavioural defences represent genomic responses to selection.

Here we leverage a dataset on Cuckoo parasitism rates at continental scale, together with genome-wide sequencing data from the Reed Warbler host (Figure 1A, Table S1). This is currently the largest available dataset with which to explore the genetic basis of antagonistic coevolution for a highly mobile avian species in which anti-parasite adaptations are predominantly behavioural in nature. Such a dataset provides us with the best available opportunity to (1) quantify variable gene flow among host populations across much of their breeding range, while (2) testing for signals of selection using genotype-environment association analyses, identifying SNP (single nucleotide polymorphism) loci associating with variation in Cuckoo parasitism rates across Europe (and controlling for evolution underpinned by variation in abiotic environmental differences). In recent years, there has been increasing recognition of the value of landscape genomics across different fields, suggesting that such an approach is worthwhile for questions involving biotic data (15, 16), particularly when signatures of adaptation might be subtle and otherwise missed by methods such as genome-wide association scans (17). Furthermore, genotype-environment association analyses enable us to take into account abiotic variables which could modulate responses to the biotic selection agent of interest (i.e. ‘landscape community genomics’ (18)). We avoid explicitly comparing the expression of behavioural defences (e.g. rate at which host parents reject foreign eggs) with genomic variation for two key reasons. Firstly, although previous studies found an association between an allele at microsatellite locus Ase64 and egg rejection behaviour in Magpies (*Pica pica*) parasitised by Great Spotted Cuckoos (*Clamator glandarius*) (19, 20), such an association was not detected in Great Reed Warblers (*Acrocephalus arundinaceus*) parasitised by Cuckoos (21), revealing the limitations of a behavioural candidate gene approach in detecting selection in the wild (7). More importantly, although behavioural data exists for some of our sampled sites, it does not take the possibility of cryptic plasticity into account (to assess this would require experimental alteration of Cuckoo activity). Instead, by considering variation in the local intensity of selection (i.e. site-level parasitism risk; Figure 1B), and by controlling for variation in abiotic environmental variables (Figure 1C), our use of genotype-environment association analyses allows us to investigate genomic evidence for a geographic mosaic of coevolution in a host-cuckoo system.

**Figure 1.**
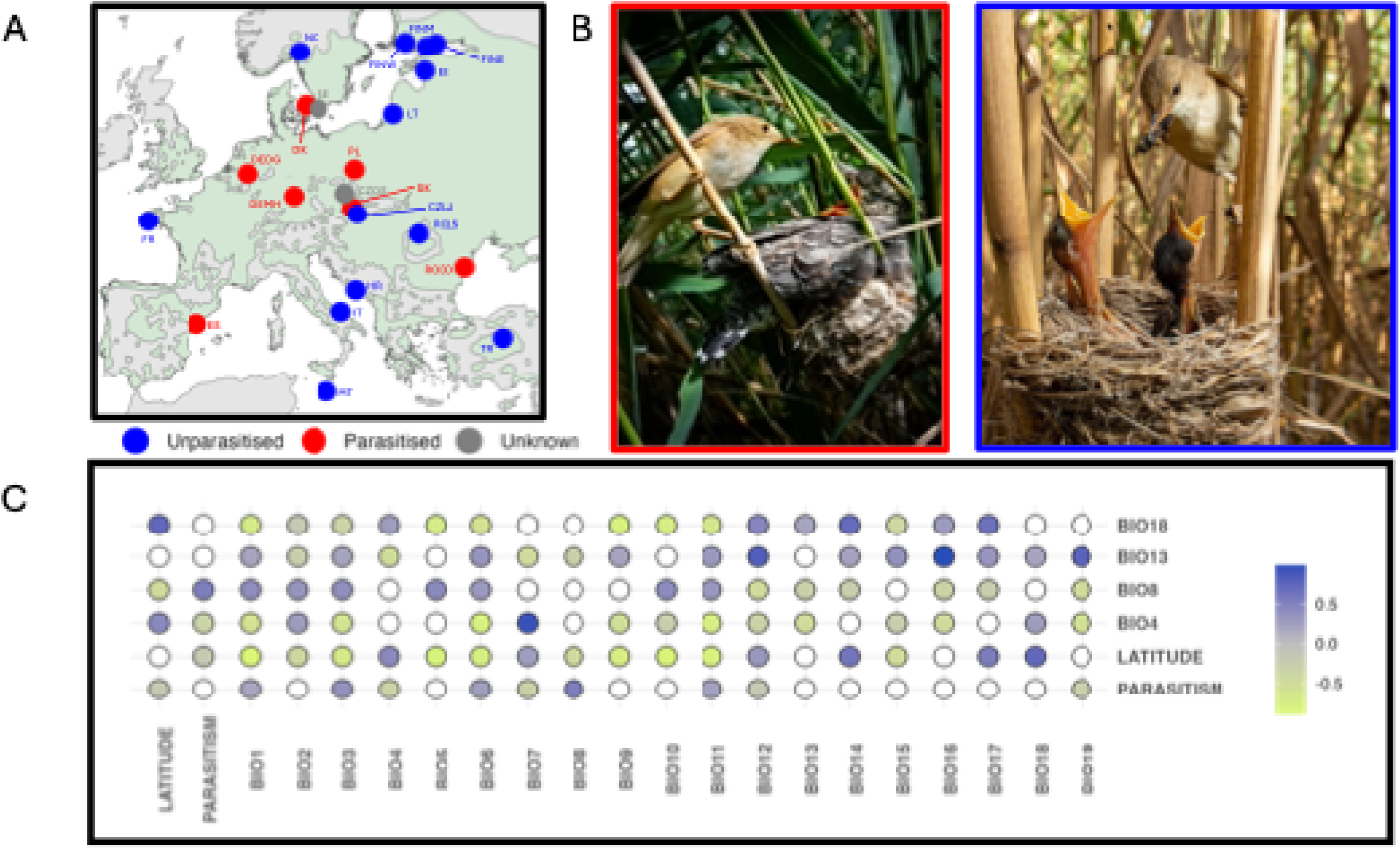
Variation in brood parasitism risk for Reed Warblers: (A) We used DNA samples from Reed Warblers at 22 sites (location codes indicated, see Table S1) across their European breeding range (coloured in light green, Birdlife International (2025)). These sites were either parasitised by Cuckoos (red points) or not parasitised (blue points). Grey points represent sites for which we currently lack information about rates of brood parasitism. (B) Brood parasitism should exert strong selection, Left – a parasitised Reed Warbler gains no reproductive success from fledging a Cuckoo chick. Right – an unparasitised Reed Warbler nest with a parent feeding its own young (Photographs by Deryk Tolman). (C) Pairwise correlations among focal environmental variables (on vertical) with other available BIOCLIM variables. Non-significant correlations (p < 0.05) are whited out. See Figure S4 for pairwise correlations amongst all BIOCLIM variables and parasitism risk.

## Methods

### Data collection

The Reed Warbler is a migratory passerine that overwinters in sub-Saharan Africa but breeds in reedbeds and reed-fringed waterways across much of Europe. Its breeding range is expanding to the north (24) and possibly, in certain regions, to the south (45). Whilst the Cuckoo is widespread, the gens laying eggs which mimic those of the Reed Warbler occurs mainly within the core of the Reed Warblers’ European breeding range. Nevertheless, even within this region parasitism varies, with Reed Warblers at sites only kilometres apart experiencing parasitism rates of between 0 and 20% of nests (46). We used sampling sites established in previous studies where Reed Warblers were caught during the breeding season and blood sampled (22-24, 36, 45). These sites formed a latitudinal and longitudinal gradient across much of Europe (Figure 1), and included locations with known range expansion histories, reducing the impact of other environmental, genetic, or demographic effects (e.g. allele surfing) on our inferences.

We collated spatial data on brood parasitism rate, making use of a combination of existing literature (39) and information provided by academic and citizen scientist collaborators (Table S1). All ‘parasitised’ sites had a parasitism rate (i.e. percentage of parasitised nests) of > 5%. All ‘unparasitised’ sites had a parasitism rate of 0% during the period in which the samples were collected (although parasitism may occur very rarely – e.g. in Finland’s Helsinki region, which hosts an ‘unparasitised’ Reed Warbler population, two nests were found to be parasitised by a (probably single) Cuckoo in 2023, out of 644 nests which reached egg stage across eight years of monitoring (pers. obs. RT). For two sampling sites, parasitism status was unknown (Figure 1A).

### Genomic data generation and pre-processing

Genomic DNA extraction and RAD-sequencing was carried out as detailed in prior work (22, 24). Briefly, DNA was extracted with QIAGEN DNeasy® Blood & Tissue Kits (Qiagen, California, U.S.A) following the manufacturer-provided protocol (24). After assessing the concentration and purity of each DNA extract, we normalised sample concentrations. Single-digest, single-end RAD-seq libraries were prepared at Floragenex (Oregon, U.S.A.). After digestion with the SbfI enzyme, sequencing was carried out on an Illumina HiSeq® 2000 platform. We generated and filtered two genotype files, one for use in population genetic structure and gene flow analyses, and another for selection detection using genotype-environment analyses (GEA). Both files included both autosomes and sex chromosomes. When selecting the samples for both datasets, we scanned for pairs of individuals with high relatedness, but none of the pairs of individuals had relatedness of > 0.1, and therefore no individuals were excluded. To calculate this information, we used VCFtools v0.1.17 (47) --relatedness. We split Finnish samples into three groups called ‘West’, ‘Middle’, and ‘East’ Finland, corresponding to three discrete sampling regions across the south coast of the country. The subsequent ‘Dataset 1’ included 190 individuals from 22 sampling sites across Europe. For the genotype file used in GEA pipelines, ‘Dataset 2’, we used a subset of the individuals from Dataset 1, excluding sampling sites for which there was no brood parasitism data (Sweden and one of the Czech Republic sites) and splitting the Finnish samples in the same way as in Dataset 1. Dataset 2 included 179 individuals from 20 sampling sites. Full details for the genomic datasets are available in Table S1.

For both Dataset 1 and Dataset 2, we used STACKS v2.65 (48) to sort, filter, and demultiplex reads with the *process_radtags* script. We aligned reads to the bAcrSci1 reference genome (31) with bwa v0.7.17 (49). We used SAMTOOLS v1.18 to sort files (50). Following this, we used the *ref_map*.*pl* pipeline to build the initial catalogue to be used as an input for *populations*. Using VCFtools, we filtered by depth, retaining sites with mean depth of more than or equal to 10, and less than or equal to 100. We then assessed sites for Hardy-Weinberg Equilibrium (using an exact test --hwe) and filtered to exclude SNPs out of HWE in all populations with more than ten individuals. Using *populations*, we selected sites where all individuals were represented (-R 1), where there was a minimum minor allele count of 3 (--min-mac 3), and where there was a maximum observed heterozygosity of 0.75 (--max-obs-het 0.75). We restricted analysis to the first SNP of each locus (--write-single-snp). For Dataset 1, we used PLINK v2.00(51) to detect SNP pairs with linkage disequilibrium of 0.5, and then exclude one of them (--indep-pairwise 50 5 0.5), retaining 13,223 SNPs (‘Dataset 1’) for our analyses. For Dataset 2, we did not filter for linkage disequilibrium as this would risk removing parts of the genome under strong selection. Hence, we retained 13,349 SNPs.

### Population genetic structure and gene flow analyses

We first carried out PCA with PLINK v2.00 (51), as well as ADMIXTURE v1.3.0 (52), ranging the number of assumed populations between 1 and 22 (i.e. the total number of sampling sites). We then examined gene flow among sampling sites. To do so, we used EEMS v0.0.0.9, which makes use of geo-referenced genetic samples to detect regions of higher- and lower-than-average gene flow and has been simulated to accurately recover effective migration surfaces (visualised gene flow) in scenarios with complicated histories of gene flow (53). We first converted the Dataset 1 VCF into PLINK input files using the –make-bed command. We then used the bed2diffs_script to convert these into a matrix of average pairwise genetic dissimilarities, which was used as the input file for EEMS. Following this, we carried out 8 runs each with an MCMC chain of 10,000,000 iterations, with a 20,000-iteration burn-in and 1800 thinning iterations. The number of geographic demes was set to 100, 300, 500, 700, 900, 1100, 1300 and 1500, and runs were combined using rEEMSplots in R. Data is presented on a log_10_ scale, where an increase of one represents a tenfold increase in migration rate relative to average levels. We also used VCFtools to output mean pairwise F_ST_ values for each population pair.

### Genotype-environment analysis

To explore genotype-environment associations we used three GEA analytical frameworks: univariate latent factor mixed modelling (LFMM) (54), multivariate redundancy analysis (RDA) and partial redundancy analysis (pRDA) (55), and the F_ST_-based method BayeScEnv (56). We used the vcf2geno command from the R package *LEA* (57) to convert the relevant VCF files into suitable input files. Finally, we estimated additive polygenic scores (58) to determine the cumulative signal of selection across candidate loci identified across all GEA methods. Parasitism rate tends to be highest in the centre of Europe (59). To summarise climatic variation, we downloaded the 19 BioClim climate variables (60). To avoid high correlations amongst the variables, we calculated Pearson’s correlation coefficient in R and selected four BioClim variables which had correlation coefficients of < 0.7 across our sites. These were temperature seasonality (BIO4), mean temperature of the wettest quarter (BIO8), precipitation of the wettest month (BIO13), and precipitation seasonality (BIO15). BIO15 was included as it was identified to potentially contribute to genetic variation in European populations of the Reed Warbler in prior work (22). BIO8 was included as it had the highest (albeit still moderate at ~0.5, see Figure S4) correlation with parasitism rate, allowing us to test for potential confounding effects. Measures of precipitation were included given the significance of rainfall in determining water level and therefore reedbed growth (i.e. Reed Warbler habitat availability and quality). We also included latitude in our models, to account for neutral geographical variation. Both LFMM and RDA/pRDA were run with parasitism rate, climatic variables, and latitude together. BayeScEnv was run individually for each variable.

We carried out LFMM with the *lfmm* v2 package in R (61). This uses an MCMC algorithm to explore associations between allele frequency and environmental variables. We included parasitism status, our four focal BioClim variables, and latitude as our environmental variables. We then ran LFMM using *lfmm_ridge* to estimate the model, and *lfmm_test* to assess the significance of the SNPs. The number of latent factors was chosen based on our population genetic structure analyses (allowing us to control for neutral genomic processes e.g. due to colonisation history). K = 1 was supported by the PCA, ADMIXTURE, and F_ST_ results, which all inferred limited differentiation across Europe. We then calculated the genomic inflation factor (GIF) (62). We confirmed that GIF values were close to 1, demonstrating that population structure was adequately controlled for. LFMM then adjusts the p-values by the GIF values, and we applied the Benjamini-Hochberg procedure (63) to tolerate a false discovery rate of 10%, to return significant candidate loci.

Following LFMM, we carried out RDA, which models relationships between environmental predictors and genomic variation (i.e. the SNPs). RDA is a constrained ordination and has been demonstrated to outperform other GEA analyses when selection acts over multiple loci (55). We carried out RDA using the *vegan* R package’s *rda* function. We calculated adjusted R^2^ (based on the number of predictors), before assessing variance explained by each eigenvalue. We assessed the RDA model for significance using F-statistics via an Analysis of Variance (ANOVA) with 999 permutations. We assessed both the full model and each constrained axis. We also assessed the Variance Inflation Factors of each predictor variable, confirming that they are below ten, and therefore that collinearity amongst predictors is not a significant problem. We identified candidate SNPs on significant constrained axes based on their loadings in ordination space, using a cut-off of ±3 SD, and removed duplicate candidates. We repeated this in a ‘partial RDA’ context, which accounts for population genomic structure. Applying both RDA approaches provides a more holistic assessment of SNPs under selection according to variation in levels of brood parasitism.

Finally, we used BayeScEnv v1.1 (56). BayeScEnv does not directly assess associations between allele frequencies and environmental variables, but rather environmental and genetic differentiation (i.e. F_ST_), providing an additional GEA test. We used VCFR (64) to convert the VCF file to a genind object (of the adegenet R package (65)), and then used hierfstat (66) to convert this into an input file. We divided parasitism rate at each site by the standard deviation, as recommended by the BayeScEnv authors (56). We then ran BayeScEnv with twenty pilot runs of length 5000, a thinning interval size of ten, 5000 outputted iterations, a burn-in length of 50,000, prior probability against the neutral model of 0.5, and prior preference for the locus-specific effect model of 0.5. We identified SNPs with a posterior error probability of < 0.01 as candidate loci. We repeated BayeScEnv using the four Bioclim variables and latitude as the focal environmental variable (zero-centered and scaled by the standard deviation), to identify any common candidate loci and therefore to test for the confounding impact of abiotic variation. We carried out three additional runs for both LFMM and BayeScEnv, generating random parasitism rate values between 0 and 25 (to be biologically realistic). This randomisation approach identified no false positive SNPs for either of the tested GEA methods, providing support for the validity of any candidates identified using the true parasitism values. RDA and pRDA determine candidate SNPs based on SNP loadings ±3 SD from the mean, and so the validity of (p)RDA candidates was assessed instead through comparison with the LFMM and BayeScEnv results.

We filtered the annotation file for the bAcrSci1 Reed Warbler genome (31) to identify candidate genes from our outlier SNPs (i.e. SNPs that fall within a gene in the annotation file). Then, we (1) performed Gene Ontology Enrichment Analysis (geneontology.org, Panther v18.0), with *Gallus gallus* (the most complete avian genome annotation) as the reference set to identify whether genes associated with specific biological processes were overrepresented, and (2) delineated the cumulative impact of selection at the individual level, by calculating per-individual additive polygenic scores (58) (estimates of the cumulative signal of selection across candidate loci; referred to as ‘polygenic scores’). Polygenic scores were calculated by computing the Pearson correlation coefficient between minor allele frequency and parasitism rate across sampling sites, to estimate the direction of the association, then extracting individual-level genotypes for the candidate SNPs in 012 format. We then aligned the SNP effects with the direction of environmental association by recoding genotypes at loci with negative correlations (to ensure that, for all SNPs, higher genotypic values reflected a positive relationship with parasitism). Finally, we calculated polygenic scores per individual as the unweighted sum of the recoded genotype values across all of our candidate loci. We assessed the correlation between the polygenic scores and parasitism rate to determine the extent to which the candidate alleles vary with parasitism. We tested linear and quadratic models and selected the model with the lowest Bayesian information criterion (BIC) value. We also assessed the correlation of polygenic scores with the bioclimatic variables and latitude, again testing both linear and quadratic models.

## Results

### High gene flow with genetic differentiation

Principal component analysis (PCA), with our dataset of Reed Warblers sampled at 22 locations across Europe, revealed that genetic differentiation identified previously along a north-south gradient (PC1) showed no discrete groupings according to parasitism status (Figure 2A) or geography (Figure S1). This corroborated pairwise F_ST_ results (Table S2) and previous studies of population structure in this species (22-24). ADMIXTURE analysis highlighted similar patterns (Figure S2). Although K = 1 was best supported, higher K values depicted some population structure, with K = 2 splitting birds latitudinally (24), characterising birds from within the range core as genetically admixed (Figure S2). There was no evidence of a split according to parasitism status (Figure S2). We then used EEMS analysis to test whether levels of gene flow would allow for a geographic mosaic of coevolution (i.e. there is neither panmixia, nor isolated populations scattered across the landscape). There was greater connectivity (higher gene flow, blue areas in Figure 2A) across the core of the continent, flanked by areas of lower gene flow in both the north and south (orange areas in Figure 2A). Although we currently lack a map of brood parasitism at adequate resolution to compare directly to this variation in gene flow, it appeared that gene flow was higher across the core of the Reed Warbler’s European breeding range, where parasitism by Cuckoos is also more common (Figure 2B). Bands of higher gene flow across longitudes, and patches of reduced gene flow latitudinally, could therefore impact connectivity of parasitised and unparasitised areas, affecting how host defence traits are distributed among populations across the breeding range (11).

**Figure 2.**
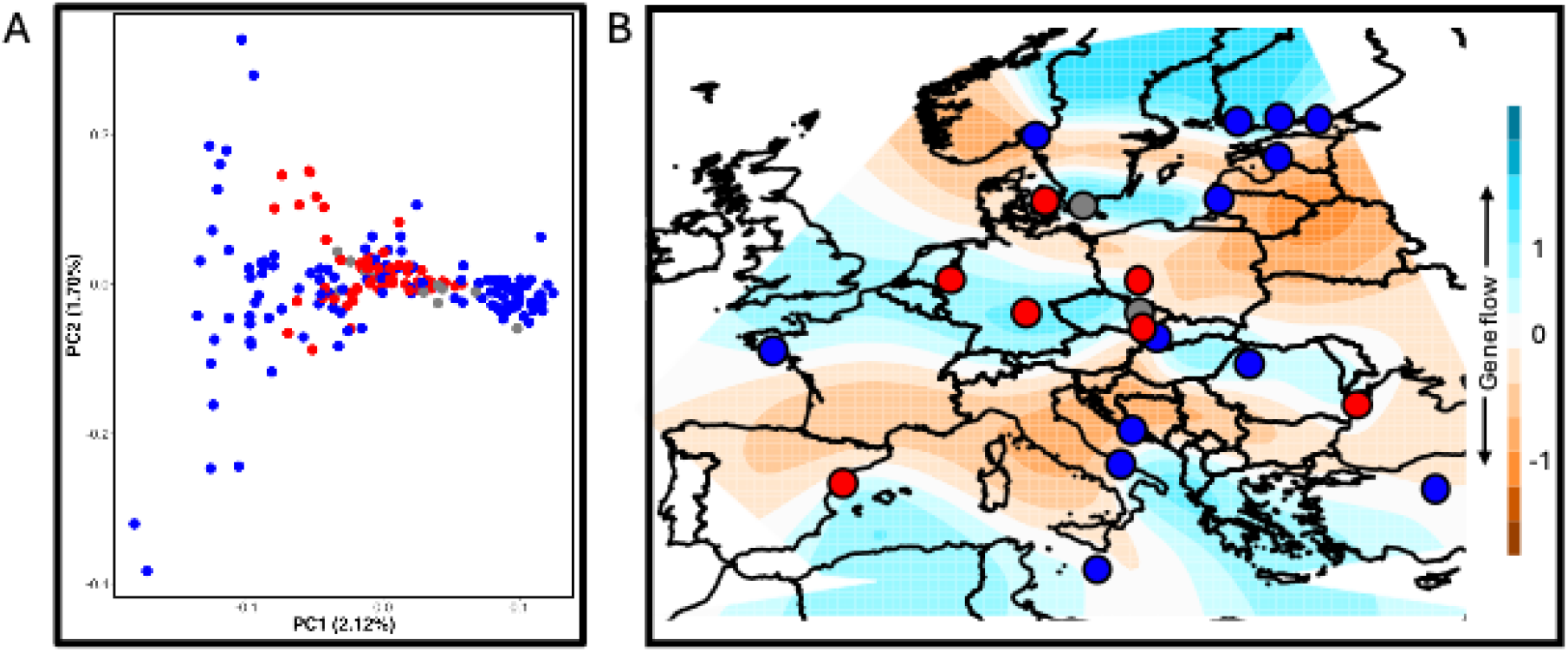
Population genetic structure of Reed Warblers according to parasitism: (A) Principal Component Analysis highlights shallow population structure without discrete groupings (variance explained by PC1 and PC2 shown in parentheses). Individuals are coloured according to parasitism status (blue = not parasitised, red = parasitised, grey = parasitism status unknown). See Figure S1 for plot by geographical locations. (B) Effective migration surfaces (EEMS) show areas with reduced gene flow (orange regions) and areas with higher gene flow (blue regions). The colour gradient represents effective migration rate on log10 scale, relative to overall migration. Gene flow is higher in the centre of Europe where parasitism is more common (sampling locations are indicated by same colour coding as in Figure 1A), and lower in the Mediterranean and Baltic regions.

### Genetic variation according to local brood parasitism rate

We used three different types of genotype-environment analysis (GEA) to investigate genetic variation associated with the local intensity of selection from brood parasitism. First, univariate latent factor mixed modelling (LFMM) identified 12 candidate SNPs associated with parasitism rate (Figure 3A, Table S3) and 145 SNPs associated with the tested abiotic environmental variables (i.e. latitude or one of four climatic variables, Figure 1C). Whilst there was some overlap of SNPs amongst our five environmental variables, the candidate ‘parasitism SNPs’ were not associated with any of these other variables. We then made use of redundancy analysis (RDA) which is a multivariate approach that combines ordination with regression of the loadings and is less prone to detecting false positives than LFMM, especially when selection acts over multiple loci (25). The global RDA model was significant to p = 0.001 with limited impact of collinearity (variance inflation factors (VIF) all < 5) and the first five axes explained 86% of the variation explained by the global model (0.89% of the total genetic variation, adjusted R^2^; note that variance explained is typically low in RDA models assessing genome-wide impacts of environmental variation). Outlier detection identified 37 SNPs associated with parasitism, 87 with latitude, and 68 with the climatic variables (Figure 3B). These SNPs were then largely confirmed by a partial RDA (pRDA), which controls for spatial factors (i.e. represents a proxy for population genetic structure) (global model significant p = 0.001, 0.44% of total genetic variance). Here, the first four axes were significant to p = 0.001 (capturing 70.06% of explained variation, Figure S3), and in outlier detection we identified 35 SNPs associated with parasitism (29 of these SNPs were identified by both RDA and pRDA, Figure 3C), 82 with latitude, and 73 with the climatic variables. Whilst the detection of SNPs associated with parasitism using RDA/pRDA may be somewhat confounded by bioclimatic variation (Figure 3B), 9 of the 12 candidate SNPs identified by LFMM were also identified by RDA and pRDA. LFMM tests each biotic and abiotic variable in a univariate (per-SNP) manner and did not identify overlap among SNPs associated with parasitism and other environmental variables, strengthening support for a direct role of parasitism risk in determining this genetic variation. Limited differences between the RDA and pRDA approaches also suggests a limited impact of population genetic structure on our results. Finally, we used a Bayesian GEA method, BayeScEnv, which assesses F_ST_-based differentiation in relation environmental variables, and has the advantage of allowing for a more diverse range of relationships between allele frequencies and the environment when compared to other methods. This method detected two further candidate parasitism SNPs. One of these loci was one which was also identified by RDA. Of the environmental variables tested with BayeScEnv, only one SNP was identified as a candidate (associated with precipitation of the wettest month). It is possible that BayeScEnv detected fewer SNPs under selection in this case study compared to the other methods because of very low between-population F_ST_ in European Reed Warblers (Table S2).

**Figure 3.**
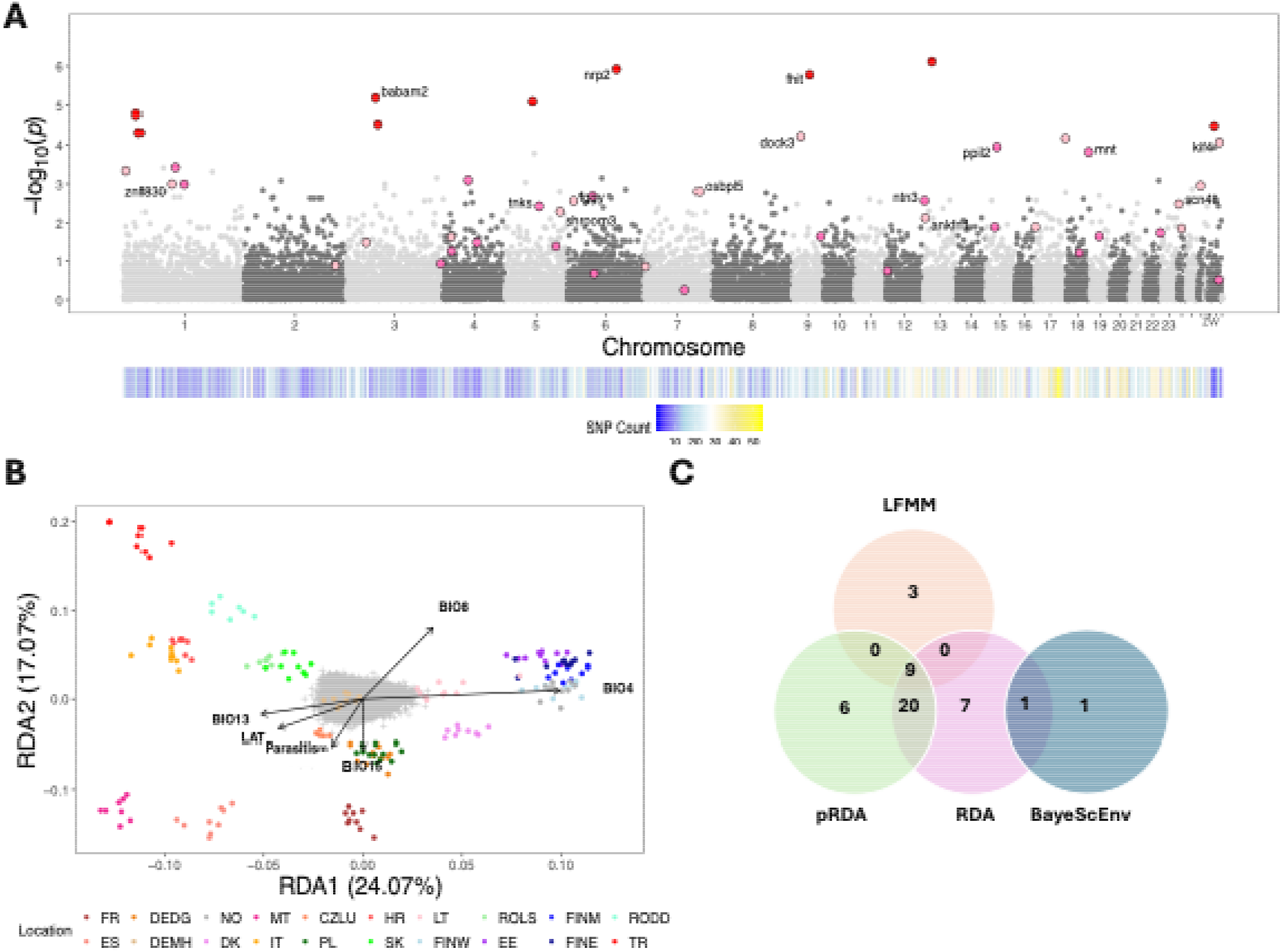
Genotype–environment association with brood parasitism: (A) Manhattan plot showing genotype–parasitism associations, estimated with LFMM, across the genome. Dots indicate the candidate SNPs identified across LFMM, RDA, pRDA, and/or BayeScEnv. Red dots are candidates identified by three GEA methods. Hot pink dots are candidates identified by two methods. Pale pink dots are candidates identified by a single method. The top ten candidates (according to LFMM) associated with known genes are annotated. Full gene names are reported in Table S3. SNP density is presented underneath as the number of a SNPs in each 1 Mb window. (B) Redundancy analysis (RDA) using 13,349 SNPs, depicting the first two axes and using the following variables: parasitism, latitude, BIO4 (temperature seasonality), BIO8 (mean temperature of wettest quarter), BIO13 (precipitation of wettest month), and BIO15 (precipitation seasonality). SNP/individual scores (coloured by location) are scaled by the square root of the eigenvalues to give appropriate weight to the axes based on their contribution to the model. SNP genotypes are represented as grey crosses, and the direction of variation in each constraining variable is indicated by black arrows (arrow length represents correlation strength). Percentages on axes indicate the proportion of variance explained. See Figure S3 for partial RDA. (C) A Venn diagram showing number of SNPs identified as varying with local brood parasitism rate by each of the different genotype-environment association methods.

Our combined approaches (LFMM, RDA/pRDA, and BayeScEnv) detected 47 ‘parasitism’ candidate SNPs distributed across the genome (Figure 3C) that overall did not correlate strongly with the other environmental variables we included. This shows that hosts have genomic variation associated with varying brood parasitism pressure and provides some evidence for a genetic basis to the hosts’ response to brood parasitism. Gene ontology analysis (using the 26 genes which were identified from our list of candidate SNPs) identified blastoderm segregation as the only process for which candidate genes were overrepresented among our candidates. As presumed adaptive responses to cuckoos by hosts are largely expressed through plastic behavioural traits (26), which are in turn expected to be highly polygenic (27), we then computed an additive polygenic score to summarise the impact of parasitism across the genome. The relationship between the cumulative signal of selection (i.e. individual-level additive polygenic scores) and parasitism rate at the associated location (*n* = 179) was of marginal significance with the linear model (*F*_(1,177)_ = 2.908, adjusted *R*^*2*^ = 0.01, p = 0.0899; Figure S5). The linear model had a lower BIC score (linear model = 721.9, quadratic model = 725.7, ΔBIC = 3.77) than the quadratic model. For all models of explanatory variables other than parasitism (i.e. latitude and the four focal climate variables), the relationships with the polygenic scores were non-significant at p > 0.2. (including BIO8, which had the highest correlation with parasitism rate of all bioclimatic variables, see Figure 1C & Figure S4), reflecting the absence of a strong relationship between parasitism and the abiotic variables used here.

## Discussion

Spatial variation in the risk of brood parasitism has often been described as forming a geographic mosaic, exerting selection on host populations at different rates (28). However, without demonstrating that there is sufficient gene flow among host populations to mix traits, or that there is an underlying genetic response to this varying selection, these descriptions have been premature. Here we present two lines of evidence for a geographic mosaic of coevolution in the host of a well-studied brood parasite. Taking advantage of many years of field research, together with available genome-wide sequencing data, first we show that the level of gene flow varies within the host’s breeding range. Panmixia would inhibit the development of a geographic mosaic of coevolution, as would complete isolation of populations across the wider landscape matrix (4). In Reed Warblers, we find higher gene flow within the range core (where there are both parasitised and unparasitised sites) and lower gene flow in the north and south of our sampling area. Variation in gene flow corresponds to general patterns in the intensity of brood parasitism at continental scale, wherein parasitism rate is generally higher in the host’s breeding range core when compared to the range edges. Such geographic variation could therefore facilitate the loss, gain or maintenance of genetic variation involved in responses to Cuckoo brood parasitism. This could allow adaptation to parasitism to vary spatially within the hosts breeding range, at continental scale. Variation in parasitism rate occurs at finer scale than we have been able to sample, so future work at more local or regional geographic scales would be valuable.

Secondly, while plasticity in behaviour poses a challenge in quantifying geographic mosaics of coevolution (29), by investigating correlations with parasitism rate we identify multiple genomic regions which are putatively under selection. Our result supports the existence of a polygenic genomic basis to hosts’ behavioural responses (i.e. adaptations to defend their reproductive success from brood parasites), suggesting that known continental-wide variation in behavioural traits may not be purely plastic (although note that putative ‘plasticity genes’ might play an important role in facilitating diverse anti-parasite responses (30)). This provides novel support for a major assumption of the GMTC in this system – that there is a genetic basis to traits involved in antagonistic interactions. Reed Warblers are known to adjust their likelihood of rejecting foreign eggs in their nests according to local parasitism risk, with individual decisions corresponding to geographic patterns (11). These patterns are reminiscent of the relationship we found here for the strength of selection from parasitism on genomic regions. In theory, field experiments to alter the hosts’ perception of parasitism could be conducted across the continent to disentangle the role of plasticity and genetic changes in determining the responses of individual hosts towards Cuckoos. In practice however, this approach would require sampling both the behaviour and genomics of hundreds to thousands of birds to derive robust conclusions. Using a proxy for the effects of behavioural interactions, as we have done here, circumvents this problem and enables us to achieve an important first step by describing a putative continental signature of genomic adaptation to brood parasitism pressure.

While determining the functional expression of genes in wild-living avian systems is still largely out of reach, we investigated evidence for potential genes associated with host responses to brood parasitism. Using the Reed Warbler genome (31) and our outlier parasitism SNPs, we identified candidate genes (Table S3) and performed a Gene Ontology Enrichment Analysis. Although most SNPs were not associated with a known gene, some were associated with a gene previously studied in avian systems (Table S3). These included genes with neurological (*nrp2*) and immunological (*ppil2*) functions. In particular, *nrp2* may potentially relate to vocal learning in birds (32) and to memory and learning in laboratory mice (33). Reed Warblers adjust vocal mobbing behaviours and egg rejection in response to changing cues of Cuckoo activity (34, 35), and it is possible that other behaviours such as nest concealment and timing of laying (36, 37) may impact on parasitism likelihood. Unfortunately there were no SNPs in the Ase64 microsatellite locus (associated with egg rejection capability in magpies (19)), precluding us from testing whether this region is associated with brood parasitism rate in Reed Warblers. Moreover, some of the putative parasitism genes, such as *ppil2* (associated with immunological changes during avian moult (38)), could suggest that some candidate SNPs might be impacted instead by correlates of parasitism rate. A well-known biotic correlate of brood parasitism is the increase in parasitism rate with host population density (39, 40). We did not have adequate data to include this variable in our analyses, so it remains a possibility that selection may be mediated by local conspecific density, or some correlate (e.g. intraspecific competition or disease), rather than directly by brood parasitism risk. Although RAD-sequencing data is ideal to assess broad-scale genome-wide variation in relation to brood parasitism (i.e. a primary goal of this study), a lack of complete genome coverage makes it probable that genes of interest under selection are left undetected. It is also probable that, rather than having a direct mechanistic role, many of our candidate SNPs are in linkage with others that are adaptively responding to parasitism rate.

Finding support for a genomic association with parasitism requires a host species which has sufficient abiotic and biotic information available, and our results highlight a need for better understanding of the natural history of brood parasitism. Weak correlation between our polygenic scores and site level parasitism is perhaps to be expected, given the potential for non-additive dynamics among genes involved in parasitism, coupled with additional environmental correlates. Variation in polygenic scores even amongst sites with no parasitism could suggest weak selection to lose anti-Cuckoo adaptations, linked selection pressures, or gene flow. Separately, we currently lack information on parasitism rates for even some of our study locations, and higher density sampling would be useful given the small scales at which brood parasitism rate varies in this system (to better characterise variable responses to selection given varying levels of gene flow (4)). Furthermore, the impact of human activity in recent centuries on changing landscapes across Europe likely affects the distributions and abundance of the Cuckoo and its hosts. The existence of gentes, each potentially subject to differing selection pressures, adds complexity to host–Cuckoo coevolutionary mosaics. Might we expect different genomic regions to be impacted by selection in different Cuckoo gentes, due to variation in parasitism pressures across host populations? Prior genomic work with Cuckoos has provided evidence for selective benefits of blue eggs (41), and of dichromatic female plumage (13), but there has been no work combining genomics and behaviour in the parasite itself.

Our characterisation of this antagonistic coevolutionary interaction represents the most in-depth assessment of the genetic underpinnings of brood parasitism carried out to date. A similar approach – using estimates of interaction strength – could hold promise in elucidating biotic interactions more generally (7, 15, 18). For example, in tight predator–prey relationships, predation rate could be used in much the same way as we have used brood parasitism (although note that, as with brood parasitism, variables including prey population density may also confound results). Such approaches are being increasingly used as part of wider population genetic studies (as well as gene expression-based assessments (42) or experimental approaches (43) which have been deployed in controlled settings). More broadly, the application of genomic research to study behaviour in well-studied systems such as mice (44) is developing rapidly. Nevertheless, genomic approaches deserve specific attention in non-model systems in the wild, especially in tightly linked predator–prey species pairs which are known to experience cyclical demographic changes. Overall, our work shows that, even with traits which are likely to be highly polygenic or involve significant plasticity, genotype– environment analyses offer opportunities to improve our understanding of the genetic underpinnings of behaviour and biotic interactions, especially when deployed alongside a thorough grounding in the natural history and field biology of the focal species. In conclusion, our investigation represents a significant step forward in understanding the geographic mosaic of coevolution in a brood parasite – host system. We not only provide support for variable gene flow across the breeding range of the host, but also provide evidence for a putative genomic signature of selection exerted by behavioural antagonists in the wild, providing support for assumptions that have underpinned interpretations of behavioural plasticity and selection in coevolutionary systems for more than fifty years.

## Acknowledgments

We are grateful to those who supplied brood parasitism rates in prior works: Geir Rudolfsen, Inge Hafstad, Bruno Bargain, Josef Beier, David Bigas Campàs, Bernd Leisler, Péter Pap, Ričardas Patapavičius, Lucyna Hałupka, Karl Schulze-Hagen, Johannes Bang, Bernd Leisler, Jesper Madsen, Anders Pape Møller, Andrzej Dyrcz, Marcel Honza, Vytautas Jusys, Eivin Røskaft, Arne Moksnes, Daniela Campobello, Matteo Caldarella, Michela Ingaramo, Vincenzo Rizzi, and Aleksi Lehikoinen. We made use of the CSC – IT Center for Science, Finland, and thank Pierre Nouhaud, Pasi Rastas, and Rafael Morales Leguey for assistance with initial exploration of the data, plus Jonna Kulmuni, Simon Martin, Jochen Wolf, and Chris Jiggins for analytical suggestions. Blood sampling was conducted with permission from national animal ethics boards. Finnish permits were granted by the Project Authorization Board of the Regional State Administrative Agency (ESAVI/7857/2018) and the Centre for Economic Development, Transport and the Environment (VARELY/758/2018).

## Funding

Funding was provided by an Academy of Finland grant to RT (no. 333801), a HiLIFE Fellows grant to RT, a Research Council of Norway grant to BGS (no. 151641/432), and an Ella and Georg Ehrnrooth Foundation grant to KR.

## Data availability

Code and RAD-sequencing barcodes are available at: https://github.com/SmithWillJ/brood-parasitism-genomics. Raw sequence data is available at: http://www.ncbi.nlm.nih.gov/sra/PRJNA1217894.

## Notes

### Competing Interest Statement

The authors have declared no competing interest.

### Summary of Updates

Minor mix-up of a handful of samples (which occurred several years ago) was detected and corrected.

