## Supplementary Tables and Figures for "Putative signatures of genomic adaptation to Cuckoo brood parasitism in Reed Warbler hosts across Europe"

**This file includes:**

Figures S1 to S4

Tables S1 to S3

SI References

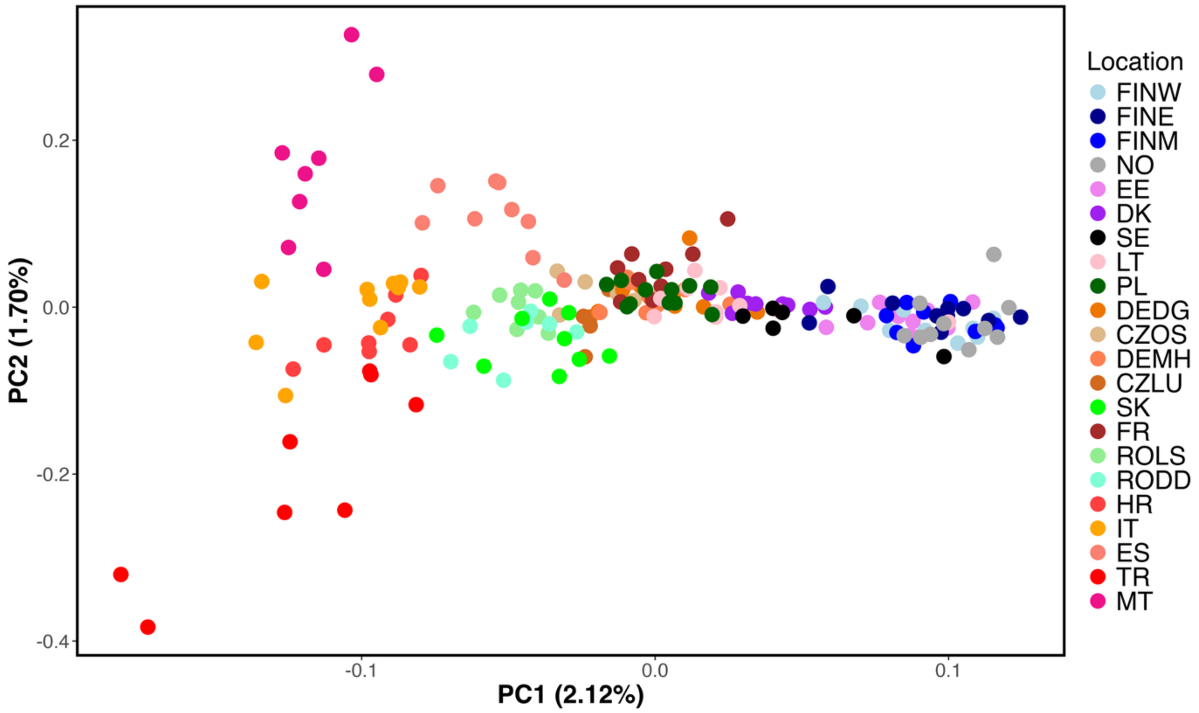


**Fig. S1.** Principal Component Analysis reveals genetic differences across the Reed Warbler range. PC1 shows a north-south gradient of genetic differences, and PC2 shows differentiation amongst southern European birds. The variance explained by PC1 and PC2 is shown in parentheses. Colours of points depict sampling regions. Sampling regions are ordered latitudinally in the figure legend. Location codes correspond to sites depicted in Figure 1A.

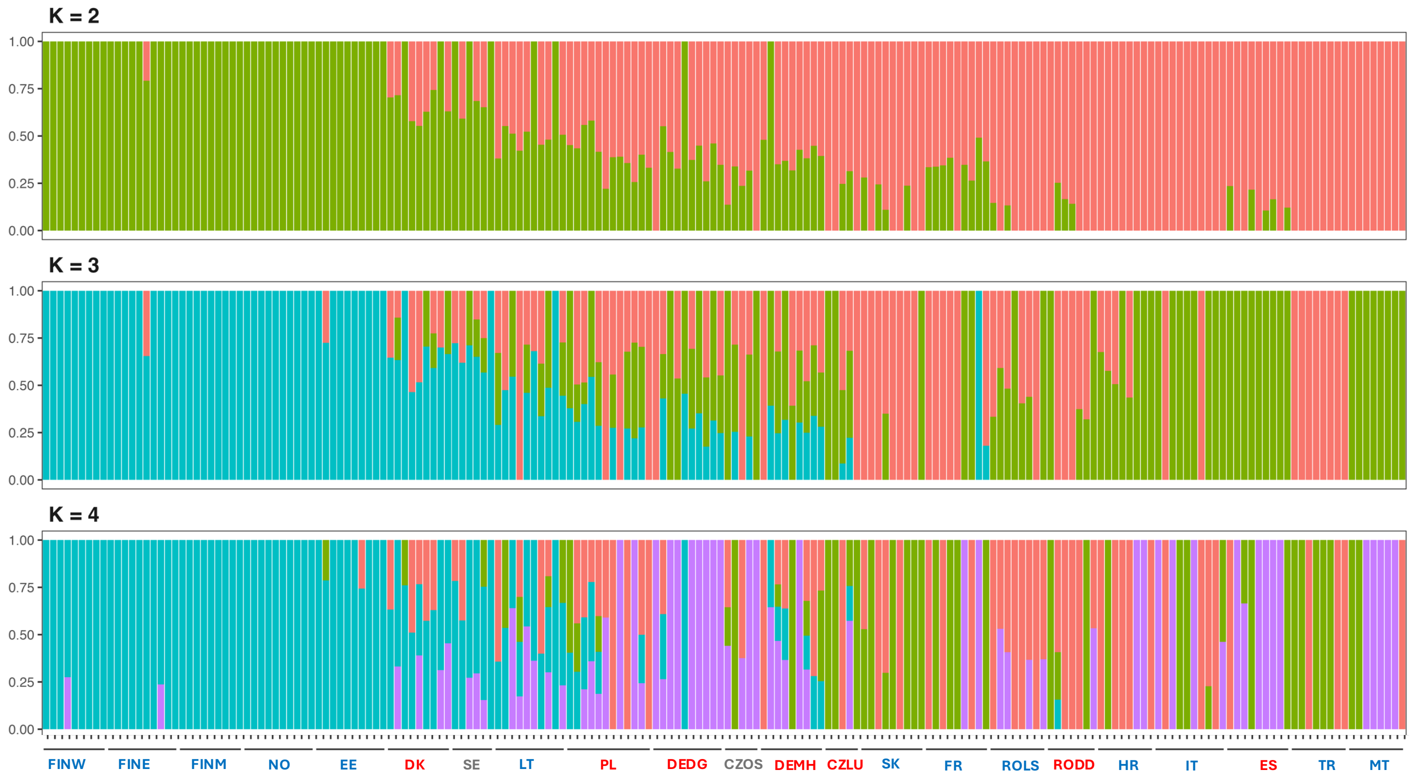


**Fig. S2.** ADMIXTURE results for 190 Reed Warbler individuals sampled from locations across Europe. Although K = 1 has the lowest CV error of the tested values, the higher K values presented here reveal some population structuring, particularly between northern and southern regions. Location names correspond to sites labelled on the map in Figure 1A. Site names are coloured according to parasitism status. Blue = unparasitised, red = parasitised, and grey = parasitism status not known. Sampling sites are ordered according to latitude.

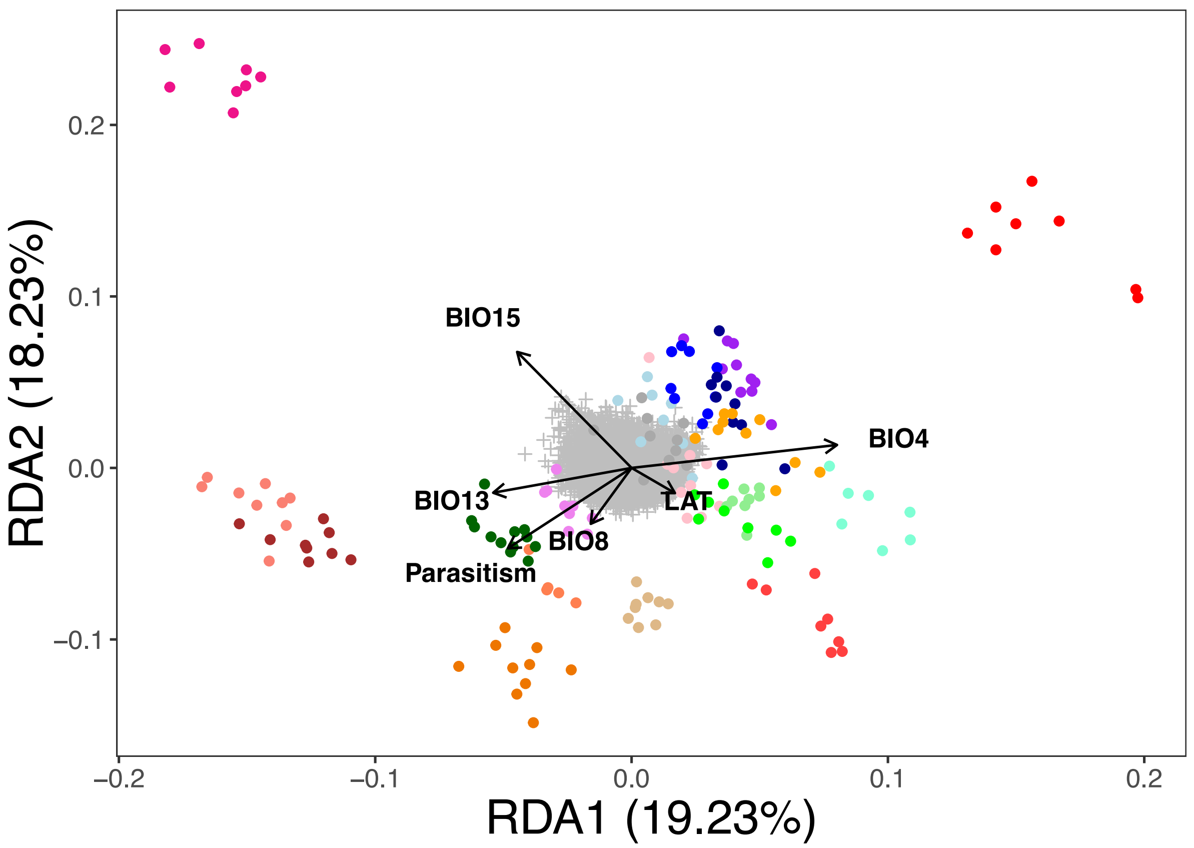


**Fig. S3.** Partial redundancy analysis (pRDA) using 13,349 SNPs, depicting the first two axes and using the following variables: parasitism, latitude, BIO4 (temperature seasonality), BIO8 (mean temperature of the wettest quarter), BIO13 (precipitation of the wettest month), and BIO15 (precipitation seasonality). SNP/individual scores are scaled by the square root of the eigenvalues to give appropriate weight to the axes based on their contribution to the model. SNP genotypes are represented as grey crosses, and the direction of variation in each constraining variable is indicated with black arrows (arrow length represents the strength of the correlation of that variable). Individuals are represented by different colours according to location (see Figure 3 for colour-coding). Percentages on the axes indicate the proportion of total variance explained.

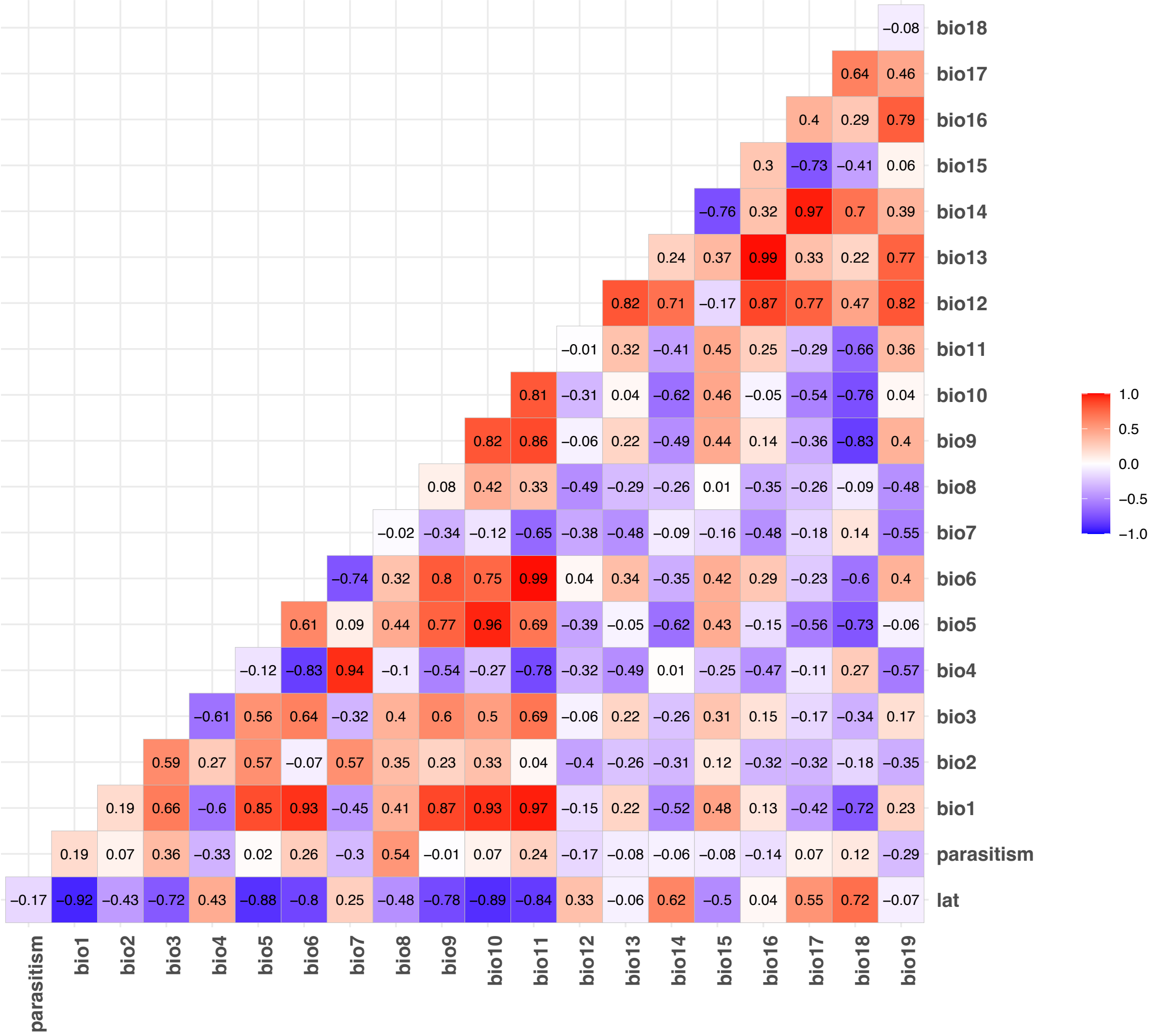


**Fig. S4.** Correlation coefficients between the bioclimatic variables, latitude, and parasitism rate. Parasitism rate is not strongly correlated with any of the environmental variables. It is moderately correlated with BIO8.

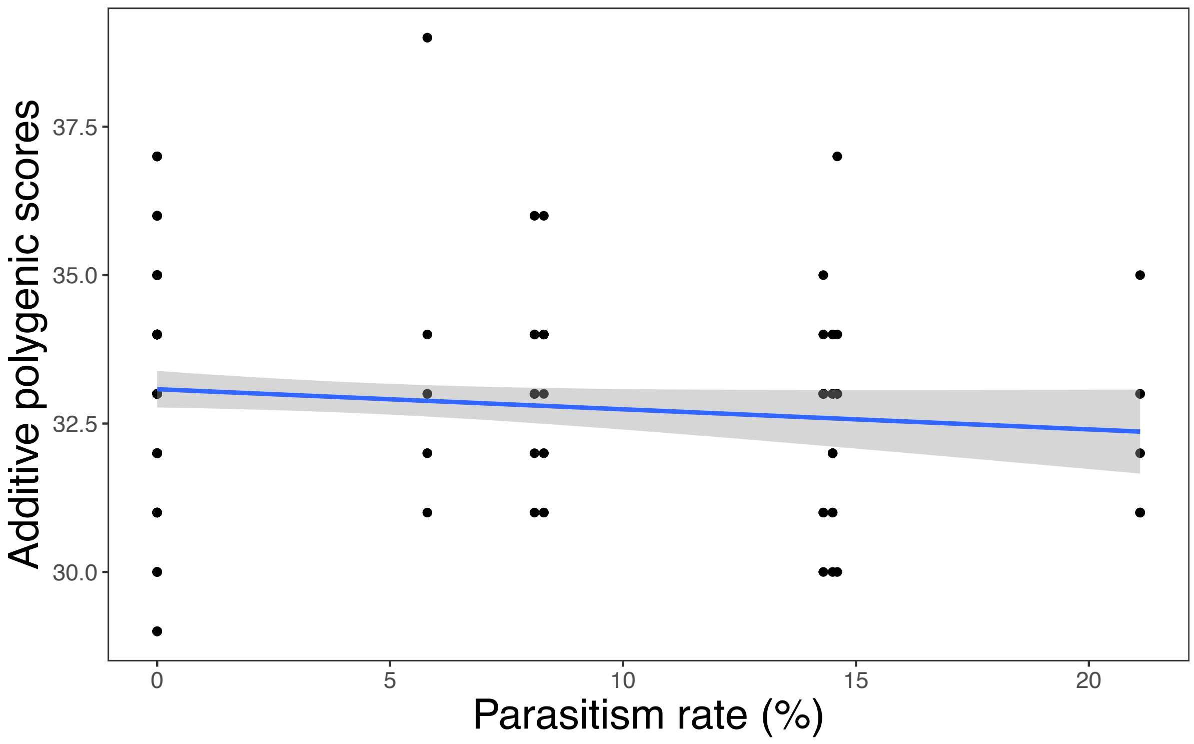


**Fig. S5.** Correlation of brood parasitism rate and individual polygenic scores (an estimate of the cumulative signal of selection across all the candidate loci). Each dot represents the polygenic score for an individual Reed Warbler and the parasitism rate at its location. The line depicts the regression line from the linear model.

**Table S1.** Review of sampling effort and the make-up of both genomic datasets used in this paper. For Finnish samples, site location is a centre of multiple sampling sites in proximity. Parasitism rate data comes from a combination of prior research, personal communications, and personal observations.

| Site name | Site code | Site location | Parasitism rate | Source for parasitism rate | Number of individuals used in 'Dataset 1' for population genetic structure (n = 190) | Number of individuals used in 'Dataset 2' for genotype-environment association analysis (n = 179) |
|---|---|---|---|---|---|---|
| Czech Republic: Lužice | CZLU | 48.852692, 17.070593 | 14.6% | Stokke et al. 2008 | 5 | 5 |
| Czech Republic: Osík | CZOS | 49.8435451, 16.2846704 | ? | N/A | 5 | 0 |
| Germany: Diergarten | DEDG | 51.233333, 6.100000 | 14.5% | Stokke et al. 2008 | 10 | 10 |
| Germany: Mohrhof | DEMH | 49.663874, 10.847465 | 8.3% | Stokke et al. 2008 | 9 | 9 |
| Denmark: Arresø | DK | 55.971545, 12.120345 | 8.1% | Stokke et al. 2008 | 9 | 9 |
| Estonia: Pärnu | EE | 58.384923, 24.533106 | 0% | *pers. obs.* BGS | 10 | 10 |
| Spain: Ebro | ES | 40.734138, 0.79174 | 21.1% | Stokke et al. 2008 | 9 | 9 |
| Finland: southeast | FINE | 60.33569, 25.69772 | 0% | *pers. obs.* RT, KR | 10 | 10 |
| Finland: middle of south | FINM | 60.14174, 24.70126 | 0% | *pers. obs.* RT, KR | 9 | 9 |
| Finland: southwest | FINW | 60.41889, 22.49918 | 0% | *pers. obs.* RT, KR | 9 | 9 |
| France: Trunvel | FR | 47.897121, -4.352747 | 0% | Stokke et al. 2008 | 9 | 9 |
| Croatia: Neretva | HR | 43.05231, 17.44803 | 0% | N/A | 8 | 8 |
| Italy: Lago Salso | IT | 41.55914, 15.87284 | 0% | *pers. obs.* KR | 10 | 10 |
| Lithuania: Ventės Ragas | LT | 55.34641, 21.240511 | 0% | *pers obs.* BGS | 10 | 10 |
| Malta: near to Xemxija | MT | 35.93749, 14.37541 | 0% | *pers. obs.* CLCS | 8 | 8 |
| Norway: Hellesjøvannet | NO | 59.73894, 11.45907 | 0% | Stokke et al. 2007 | 10 | 10 |
| Poland: Milicz | PL | 51.534002, 17.314939 | 14.3% | *pers. comm.* Lucyna Hałupka | 12 | 12 |
| Romania: Danube | RODD | 44.632193, 28.872278 | 5.8% | Stokke et al. 2008 | 7 | 7 |
| Romania: Lake Sic | ROLS | 46.966139, 23.899431 | 0% | Stokke et al. 2008 | 8 | 8 |
| Sweden: Krankesjön | SE | 55.70043, 13.47564 | ? | *pers. obs.* BGS | 6 | 0 |
| Slovakia: Trnava | SK | 48.37091, 17.58332 | 0 | *pers. obs.* BGS | 9 | 9 |
| Turkey: Mogan | TR | 39.76333, 32.79434 | 0 | *pers. obs.* BGS | 8 | 8 |

**Table S2.** Mean pairwise $F_{ST}$ between all pairs of sampled populations. Negative values are reported as zero.

| | CZLU | CZOS | DEDG | DEMH | DK | EE | ES | FINE | FINM | FINW | FR | HR | IT | LT | MT | NO | PL | RODD | ROLS | SE | SK |
|---|---|---|---|---|---|---|---|---|---|---|---|---|---|---|---|---|---|---|---|---|---|
| **CZOS** | 0 | | | | | | | | | | | | | | | | | | | | |
| **DEDG** | 0.077 | 0.077 | | | | | | | | | | | | | | | | | | | |
| **DEMH** | 0.065 | 0.065 | 0.071 | | | | | | | | | | | | | | | | | | |
| **DK** | 0 | 0 | 0.006 | 0 | | | | | | | | | | | | | | | | | |
| **EE** | 0.077 | 0.077 | 0 | 0 | 0.006 | | | | | | | | | | | | | | | | |
| **ES** | 0.065 | 0.065 | 0.288 | 0.271 | 0 | 0.173 | | | | | | | | | | | | | | | |
| **FINE** | 0 | 0 | 0 | 0 | 0 | 0 | 0 | | | | | | | | | | | | | | |
| **FINM** | 0 | 0 | 0.006 | 0 | 0 | 0.006 | 0 | 0 | | | | | | | | | | | | | |
| **FINW** | 0.129 | 0 | 0.199 | 0.188 | 0.067 | 0.199 | 0.188 | 0.082 | 0.067 | | | | | | | | | | | | |
| **FR** | 0 | 0.065 | 0.006 | 0 | 0 | 0.006 | 0 | 0 | 0 | 0.188 | | | | | | | | | | | |
| **HR** | 0.052 | 0.052 | 0.015 | 0.008 | 0 | 0 | 0.127 | 0 | 0 | 0.175 | 0 | | | | | | | | | | |
| **IT** | 0 | 0 | 0 | 0 | 0 | 0 | 0 | 0 | 0 | 0.082 | 0 | 0 | | | | | | | | | |
| **LT** | 0.006 | 0 | 0.056 | 0.048 | 0 | 0.056 | 0.048 | 0 | 0 | 0.011 | 0.048 | 0.040 | 0 | | | | | | | | |
| **MT** | 0.052 | 0.052 | 0.181 | 0.163 | 0 | 0.061 | 0 | 0 | 0 | 0.175 | 0 | 0.016 | 0 | 0.040 | | | | | | | |
| **NO** | 0.077 | 0.077 | 0 | 0 | 0.006 | 0 | 0.173 | 0 | 0.006 | 0.199 | 0.006 | 0 | 0 | 0.056 | 0.061 | | | | | | |
| **PL** | 0.100 | 0.100 | 0 | 0.086 | 0.017 | 0.010 | 0.318 | 0.010 | 0.017 | 0.221 | 0.017 | 0.027 | 0.010 | 0.069 | 0.213 | 0.010 | | | | | |
| **RODD** | 0.037 | 0.037 | 0.027 | 0 | 0 | 0 | 0.233 | 0 | 0 | 0.161 | 0 | 0 | 0 | 0.030 | 0.120 | 0 | 0.042 | | | | |
| **ROLS** | 0.052 | 0.052 | 0.015 | 0.053 | 0 | 0 | 0.253 | 0 | 0 | 0.175 | 0 | 0 | 0 | 0.040 | 0.143 | 0 | 0 | 0.010 | | | |
| **SE** | 0 | 0 | 0.047 | 0.037 | 0 | 0.047 | 0.037 | 0 | 0 | 0.010 | 0.037 | 0.026 | 0 | 0 | 0.026 | 0.047 | 0.064 | 0.014 | 0.026 | | |
| **SK** | 0.065 | 0.065 | 0.048 | 0.063 | 0 | 0 | 0.271 | 0 | 0 | 0.188 | 0 | 0.008 | 0 | 0.048 | 0.163 | 0 | 0 | 0.019 | 0.008 | 0.037 | |
| **TR** | 0.052 | 0.052 | 0.015 | 0.053 | 0 | 0 | 0.253 | 0 | 0 | 0.175 | 0 | 0 | 0 | 0.040 | 0.143 | 0 | 0 | 0.010 | 0 | 0.026 | 0.008 |

**Table S3.** Candidate genes associated with parasitism. Gene identification and/or information on function was available for 26 of the 47 candidate SNPs identified as associated with parasitism rate by one or more of the genotype-environment analysis types used for this project. LFMM = latent factor mixed modelling. (p)RDA = (partial) redundancy analysis.

| **Gene** | **Citations** | **Associations reported in prior studies using avian systems** | **LFMM?** | **RDA?** | **pRDA?** | **BayeScEnv?** |
|---|---|---|---|---|---|---|
| babam2 (BRISC and BRCA1 A complex member 2) | - | - | ✓ | ✓ | ✓ | - |
| fhit (fragile histidine triad diadenosine triphosphatase) | - | - | ✓ | ✓ | ✓ | - |
| nrp2 (neuropilin 2) | (1) | Potentially involved in vocal learning (i.e. birdsong). | ✓ | ✓ | ✓ | - |
| celf2 (CUGBP Elav-like family member 2 | - | - | - | ✓ | ✓ | ✓ |
| tbx3 (T-box transcription factor 3) | (2) | Involved in limb development. | - | ✓ | - | ✓ |
| efnb1 (ephrin B1) | (3, 4) | Expression varies with heat stress in domestic chickens. Expression varies with infection status in domestic chickens. | - | ✓ | ✓ | - |
| fam167a (family with sequence similarity 167 member A) | - | - | - | ✓ | ✓ | - |
| ppil2 (peptidylprolyl isomerase-like 2) | (5) | Related to immunity in moulting domestic chickens. | - | ✓ | ✓ | - |
| tnks (tankyrase 1) | - | - | - | ✓ | ✓ | - |
| itpr1 (inositol 1,4,5-trisphosphate receptor type 1) | (6) | Expressed in quail eggs. | - | ✓ | ✓ | - |
| ntn3 (netrin 3) | (7) | Involved in stress response in chickens. | - | ✓ | ✓ | - |
| col11a1 (collagen type 11 alpha 1 chain) | (8-10) | Associated with dwarfism in domestic chickens. Upregulated in domestic duck pituitary glands. Expression in chickens varies with abdominal fat. | - | ✓ | ✓ | - |
| dlg3 (disks large homolog 3) | - | - | - | ✓ | ✓ | - |

| | | | | | | |
|---|---|---|---|---|---|---|
| rtn4rl1 (reticulon 4 receptor like 1) | (11) | Expression levels studied in the context of avian vocal learning. | - | ✓ | ✓ | - |
| mnt (MAX network transcriptional repressor) | - | - | ✓ | ✓ | - | - |
| scn4a (sodium voltage-gated channel alpha subunit 4) | (12) | Under positive selection in toxic bird species. | - | ✓ | - | - |
| ankfn1 (ankyrin repeat and fibronectin type 3 domain containing 1) | (13) | Studied in the context of feather-pecking behaviour in captive chickens. | - | ✓ | - | - |
| znf830 (zinc finger protein 830) | - | - | - | ✓ | - | - |
| fggy (FGGY carbohydrate kinase domain containing) | (14) | Associated with red plumage colouration. | - | ✓ | - | - |
| shroom3 (shroom family member 3) | - | - | - | ✓ | - | - |
| cbs (cystathionine-beta-synthase) | (15) | Developmental role in chickens. | - | - | ✓ | - |
| alk (ALK receptor tyrosine kinase) | (16) | Studied in the context of vocal rhythm in birds. | - | - | ✓ | - |
| fancm (FA complementation group M) | (17) | Widely conserved gene involved in DNA repair – has been studied in chickens. | - | - | ✓ | - |
| osbpl5 (oxysterol binding protein like 5) | (18) | Immunological role in chickens. | - | - | ✓ | - |
| dock3 (dedicator of cytokinesis 3) | - | - | ✓ | - | - | - |
| kif4 (kinesin family member 4) | - | - | ✓ | - | - | - |
